# Interaction with single-stranded DNA-binding protein modulates *Escherichia coli* RadD DNA repair activities

**DOI:** 10.1101/2022.12.05.519199

**Authors:** Miguel A. Osorio Garcia, Elizabeth A. Wood, James L. Keck, Michael M. Cox

## Abstract

The bacterial RadD enzyme is important for multiple genome maintenance pathways, including RecA DNA strand exchange and RecA-independent suppression of DNA crossover template switching. However, much remains unknown about the precise roles of RadD. One potential clue into RadD mechanisms is its direct interaction with the single-stranded DNA binding protein (SSB), which coats single-stranded DNA exposed during genome maintenance reactions in cells. Interaction with SSB stimulates the ATPase activity of RadD. To probe the mechanism and importance of RadD/SSB complex formation, we identified a pocket on RadD that is essential for binding SSB. In a mechanism shared with many other SSB-interacting proteins, RadD uses a hydrophobic pocket framed by basic residues to bind the C-terminal end of SSB. RadD variants that substitute acidic residues for basic residues in the SSB binding site impair RadD/SSB complex formation and eliminate SSB stimulation of RadD ATPase activity *in vitro*. Mutant *E. coli* strains carrying charge reversal *radD* changes display increased sensitivity to DNA damaging agents synergistically with deletions of *radA* and *recG*, although the phenotypes of the SSB-binding *radD* mutants are not as severe a full *radD* deletion. This suggests that RadD has multiple functions in the cell, with a subset requiring the interaction with SSB.

## Introduction

DNA recombination and repair are essential for genome integrity (1, 2). In bacteria, RecA-dependent recombinational repair (3–5) and RecA-independent template switching (6, 7) are important pathways that contribute to several DNA repair mechanisms. RecA-dependent pathways appear to be the primary recombinational repair pathway used by bacteria, with RecA serving as a motor protein that mediates homologous strand exchange. Branched DNA structures created by recombination require timely resolution for cell viability (8). Accordingly, several proteins aid in the resolution of RecA-dependent and -independent repair intermediates including RecG, Uup, RadA, and RadD (8–12).

The RadD protein promotes RecA-dependent strand exchange (9) and suppresses crossover events in RecA-independent template switching (13). Deletion of the *radD* gene sensitizes *E. coli* to radiation and chemical DNA damage (14). Genetic studies have identified epistatic relationships between *radD* and genes encoding branched DNA binding and remodeling enzymes such as RadA (14), Uup (13), RecG (8) in *E. coli* and *recQ* in *V. cholera* (15). These DNA repair proteins all have roles in binding and/or resolving branched DNA repair substrates. The RadD protein is a putative superfamily 2 (SF2) helicase, with eight well conserved helicase motifs (motifs 0, I, Ia, and II-VI) (16) and demonstrated binding to forked DNA structures *in vitro* (13). However, DNA unwinding activity has not been detected for RadD. RadD acceleration of RecA-mediated DNA strand exchange requires RadD ATPase function and, presumably, interaction between RadD and RecA (9). RadD ATPase activity is also important *in vivo* as *radD* ATPase mutants are sensitized to radiation damage, although not as strongly as full *radD* deletion mutants (14).

*E. coli* RadD directly interacts with the single-stranded (ss) DNA-binding protein (SSB), forming a complex that requires the presence of the C-terminal end of SSB (17). SSB binds and protects ssDNA, while simultaneously acting as a hub for DNA metabolism through direct protein-protein interactions (18). Interestingly, RadD ATPase activity is stimulated by SSB or a peptide comprising the final nine residues of SSB (SSB-Ct) whereas ATPase activity is independent of DNA (17). This behavior has not been observed with other SSB interaction partners and the potential importance of the RadD/SSB interaction *in vivo* has not been investigated.

Many questions remain regarding the function of RadD and the possible role that its physical interaction with SSB might play in cellular DNA repair. To better understand the physical basis of the RadD/SSB interaction and its importance *in vivo*, we have mapped the SSB binding site on RadD and investigated the effects of mutations that impair Rad/SSB complex formation. Three evolutionarily conserved Arg residues frame the SSB-binding pocket in RadD. The SSB-binding pocket is connected to the ATPase active site through a helix, suggesting a possible structural link between SSB binding to ATPase stimulation in RadD. Charge-reversal sequence changes to any of the Arg residues in the pocket impairs RadD interactions with SSB and obviates SSB-stimulation of RadD ATPase activity *in vitro*. Mutation of the *radD* gene in *E. coli* with any of the charge reversal mutations had no impact on their own but, when combined with a *recG* deletion mutation, they led to induction of the SOS DNA damage response and sensitization to DNA damaging agents. Thus, the interaction between RadD and SSB is important for genome maintenance in some contexts in cells.

## Results

### Putative RadD SSB-binding pocket

To examine the role of RadD complex formation with SSB *in vitro* and *in vivo*, we first sought to identify the SSB interaction site on RadD. Crystallographic and NMR studies have identified SSB binding sites for several bacterial proteins, including exonuclease I (19), RecO (20), RecQ (21), the chi subunit of DNA polymerase III (22), PriA (23), PriC (24, 25), and ribonuclease HI (26). In each case, residues from the evolutionarily conserved C-terminus of SSB (SSB-Ct: Met-Asp-Phe-Asp-Asp-Asp-Ile-Pro-Phe in *E. coli* SSB) dock onto a pocket on the surface of the interacting protein in a manner that accommodates the side chain and α-carboxyl groups from the C-terminal Phe and side chains of upstream Asp residues. Notably, sequence changes in SSB-Ct binding pockets that disrupt complex formation with SSB lead to loss of coordinated activity with SSB *in vitro* and/or phenotypic impacts *in vivo* (19–28). A possible protein interaction role for an intrinsically disordered element with SSB has also been proposed (29).

We took a modelling approach to identify regions of *E. coli* RadD that could serve as binding sites for SSB. AlphaFold2 (30, 31) was used to predict the complex formed between the final four residues of the SSB-Ct (Asp-Ile-Pro-Phe) and RadD. The top multimer solutions converged with the SSB-Ct binding to a pocket on the surface of the RecA-like motor domain 1 (RD1) of RadD (Figure 1A). The identified site comprises a hydrophobic pocket framed by basic residues, which shares strong electrostatic similarity with other structurally defined SSB-Ct interactions sites (Figure 1B) and is evolutionarily well conserved among bacterial RadDs (Figure 1C). In the top solution (Figure 1D), Arg49 from RadD forms a bifurcated salt bridge with the α-carboxyl group of the C-terminal Phe of SSB, with the Phe side chain docked within a hydrophobic pocket. Other solutions place the α-carboxyl group of the Phe between Arg21 and Arg49. A third Arg (residue 145) is near Arg 21 and Ar49, making it available to potentially interact with the α-carboxy group or side chains of the upstream Asp residues. In all solutions, the Phe side chain docks into the hydrophobic pocket produced by the side chains of Thr16, Leu17, Phe20, Leu44, Leu113, and Leu147 from RadD.

**Figure 1.**
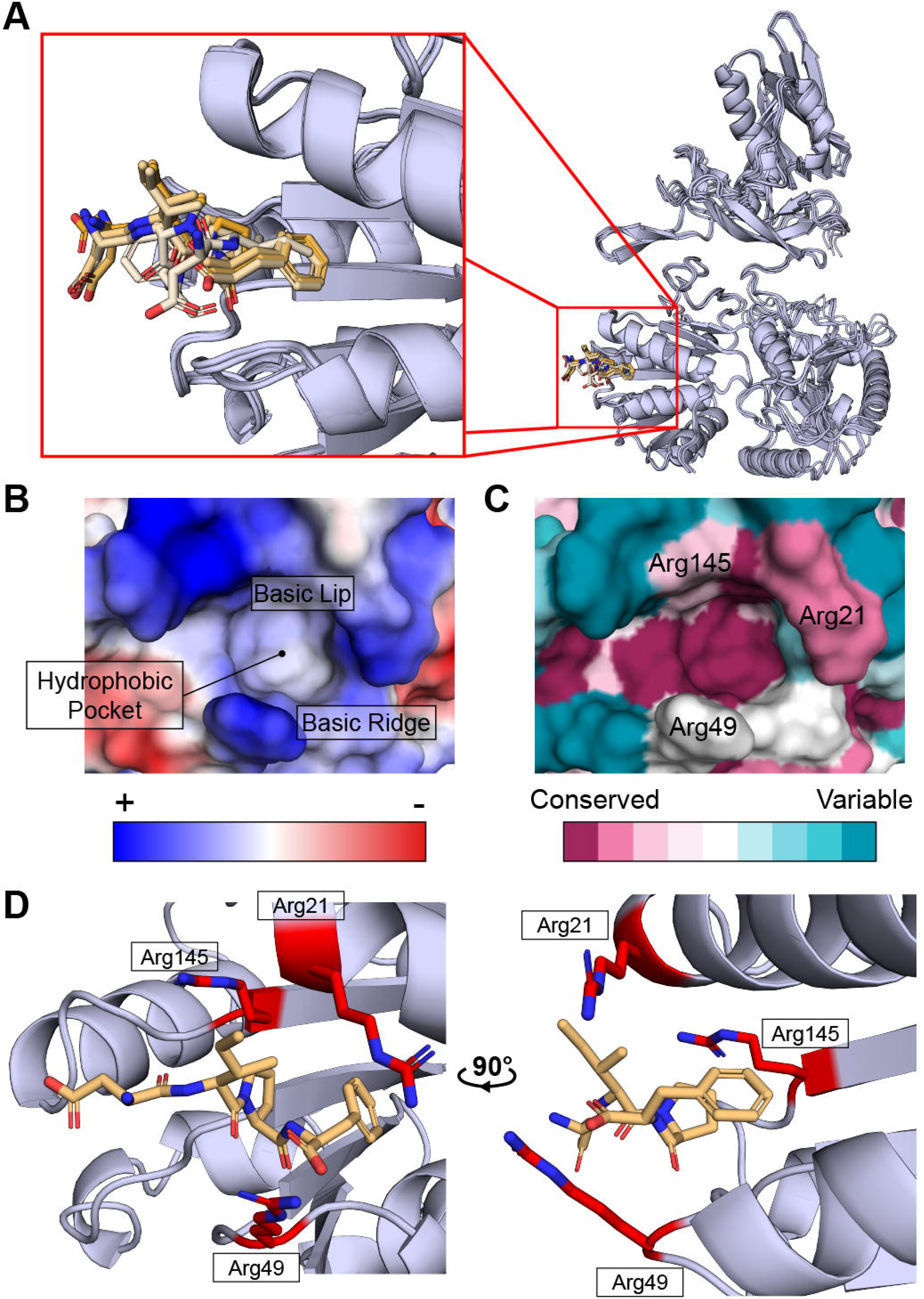
Putative RadD SSB-binding pocket A) Aligned predicted AlphaFold2 models of RadD SSB-binding pocket occupied by SSB C-terminal peptide (DIPF) in sticks. Zoomed in panel shows the top five ranked positions of the SSB-peptide on RecA-like domain 1 (RD1) colored by rank (top rank orange to bottom rank in light orange). B) Calculated vacuum electrostatics profile of putative RadD SSB-binding pocket (PDB 7R7J, (16)). Characteristic features labeled (hydrophobic pocket, basic lip and basic ridge). C) Evolutionary conservation of putative binding pocket calculated by ConSurf (52). Surface arginines responsible for charge profile of the SSB-binding pocket labeled. D) Top structure prediction of putative RadD SSB-binding pocket with SSB C-terminal peptide in light orange. Basic arginine residues outlining the pocket shown as red sticks.

### RadD SSB-binding mutants have reduced binding of SSB

To assess the putative RadD SSB-binding pocket, the codons for residues Arg21, Arg49 or Arg145 were individually mutagenized to encode for Glu (reversing their charge) in an overexpression plasmid. The variant proteins and wildtype (WT) *E. coli* RadD were then purified. The impact of the sequence changes was initially tested using a fluorescence SSB-Ct peptide binding bind assay. In this assay, the fluorescence anisotropy of an SSB-Ct peptide with an N-terminal FAM label was measured as RadD or a variant protein was titrated – binding slows rotation of the peptide, leading to an increase in fluorescence anisotropy. Consistent with prior results (17), WT RadD produced a concentration-depended increase in fluorescence anisotropy of the probe, binding with an apparent dissociation constant (K_D_) of 5.4 ± 1.3 μM (Figure 2A). In contrast, each of the RadD variants displayed severely weakened binding to the peptide probe. K_D_ values for SSB-Ct binding by the variants could not be determined, but the constants are well in excess of the highest RadD concentration tested (30 μM). These data are consistent with Arg21, Arg49, and Arg145 playing important roles in binding to the SSB-Ct as predicted by the model.

**Figure 2.**
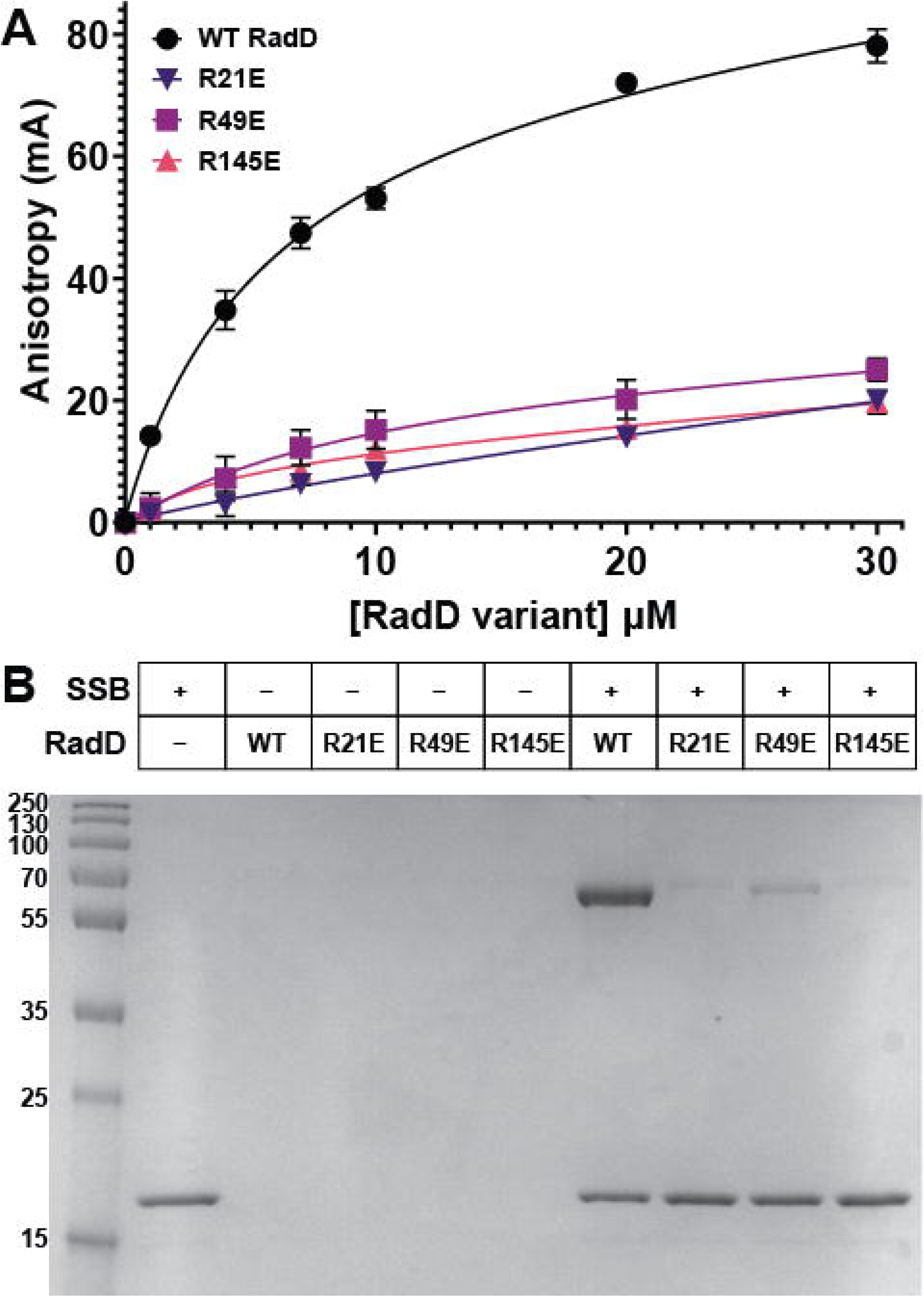
Charge-reversal changes to putative RadD SSB-binding pocket residues impairs binding to SSB. A) Fluorescence anisotropy of FAM labeled SSB C-terminal peptide WMDFDDDIPF with RadD or RadD variants. Data points are the mean of 3 independent measurements, with error bars representing mean standard error. B) Gel testing ammonium sulfate co-precipitation of RadD or RadD variants with full length SSB.

Complex formation between the RadD variants and full-length *E. coli* SSB was next tested using an ammonium sulfate co-precipitation assay (32, 33). SSB precipitates in low concentrations of ammonium sulfate under which most proteins, including RadD, remain soluble. However, formation of the RadD/SSB complex leads to RadD co-precipitation with SSB under the same conditions (17). When incubated with 150 g/L ammonium sulfate, isolated WT RadD and the RadD variants remained soluble, whereas SSB readily precipitated (Figure 2B). As observed previously, WT RadD co-precipitates with SSB, consistent with their direct interaction. In contrast, co-precipitation was greatly diminished for each of the charge-reversal RadD variants, indicating that the sequence changes strongly impaired complex formation with SSB. Together with the peptide binding results, these results show that Arg21, Arg49 and Arg145 are important for SSB binding, consistent with the RadD/SSB interface identified by modelling.

### SSB-binding pocket Arg mutants abolish SSB-specific ATPase stimulation

SF2 helicases often display DNA-dependent ATPase activities that rely on allosteric connections between DNA binding and ATPase sites (34–37). In contrast, RadD ATPase activity does not depend on DNA but instead is stimulated by SSB or SSB-Ct binding (17). Thus, interaction with SSB may directly stimulate RadD biochemical functions. The RadD ATPase active site resides in a cleft between its RD1 and RD2 domains. RD1 makes up the majority of the ATPase binding pocket, spanning SF2 helicase motifs 0, I, Ib, II, and III, whereas RD2 contains motifs IV, V and IV (16). The RadD SSB-binding pocket identified above is located on the RD1 domain surface, at a position that is quite far from the ATPase active site (Figure 3A). How SSB binding can alter ATPase activity is therefore unclear.

**Figure 3.**
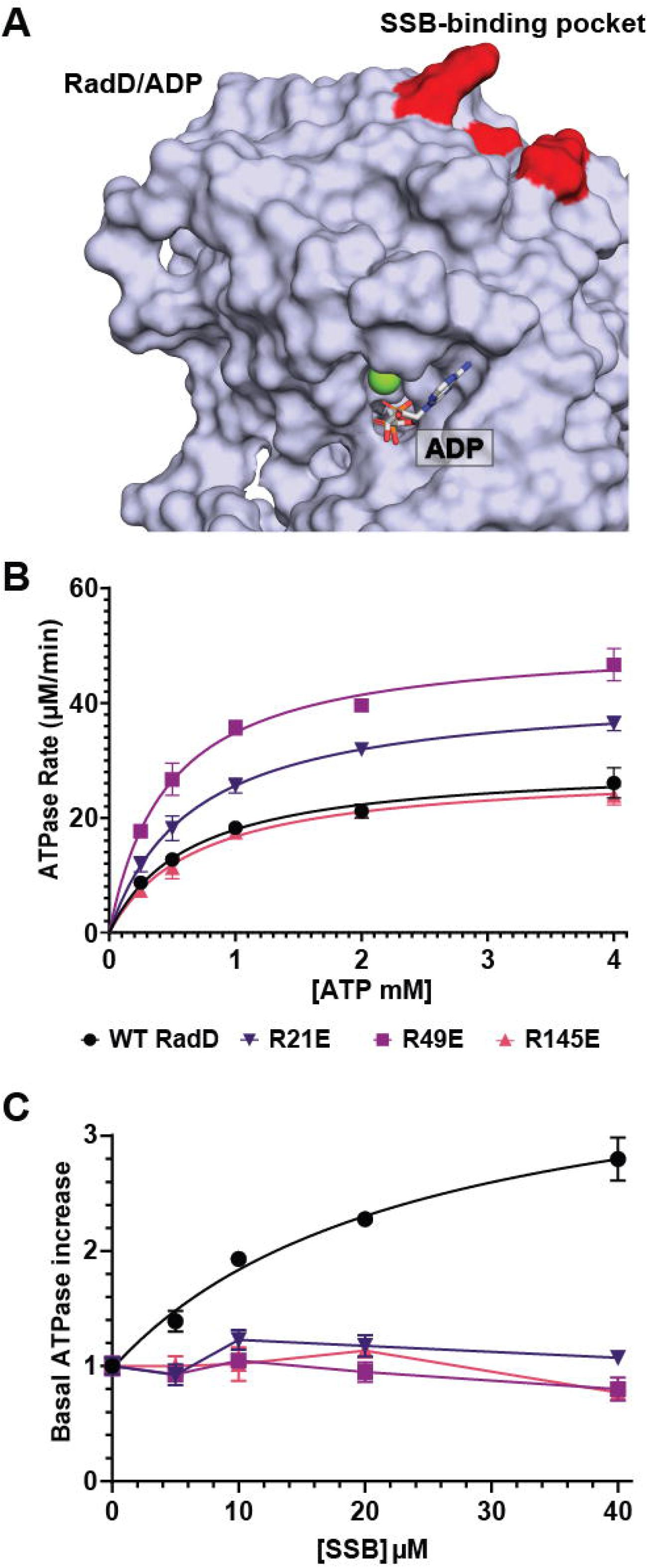
Effect of SSB-binding RadD variant sequence changes on ATPase activity A) Surface representation of RadD RecA like domain 1 (RD1) bound to ADP (PDB 7R7J, (16)) in grey. Bound ADP shown as stick in white and magnesium ion in green. Arginines important to SSB-interaction in red, located at the opposite side of the ATP binding pocket demarked by bound ADP. B) ATP-dependent ATP hydrolysis rates for 200 nM RadD or RadD variants. Data points are the mean of 3 independent measurements, with error bars representing mean standard error. C) Effect of adding SSB to 200 nM RadD or RadD variant ATPase activity. RadD ATPase induction is fit to dose response increase curve while RadD SSB-binding mutant ATPase points are connected by lines. Data points are the mean of 3 independent measurements, with error bars representing mean standard error. Rates are normalized to the V_max_ values determined in the absence of SSB.

To examine how SSB binding influences ATPase function in RadD, we measured ATP hydrolysis activity of WT RadD and the SSB site variants in the absence and presence of SSB. Steady-state ATPase kinetic parameters were measured first for WT RadD and each of the RadD variants in the absence of SSB to determine if the variant sequence changes altered basal ATPase function. WT RadD hydrolyzed ATP at a maximum rate (V_max_) of 22.5 ± 0.5 μM/min and a Michaelis constant (K_m_) of 640 ± 76 μM (Figure 3B and Table 1), consistent with prior measurements (17). The RadD charge-reversal variants each hydrolyzed ATP with K_m_ and V_max_ values that were within one-fold differences from WT RadD (Table 1). The very modest differences indicate that sequence changes to the SSB-binding site on RadD did not significantly impact basal ATPase function and that the variants are properly folded.

**Table 1.** Steady-state ATPase kinetic parameters of RadD SSB-mutants

| **Proteins** | **K<sub>M</sub> (μM)** | **V<sub>max</sub> (mM/min)** |
| --- | --- | --- |
| WT RadD | 640 ± 76 | 29.4 ± 1.13 |
| RadD R21E | 654 ± 52 | 42.3 ± 1.10 |
| RadD R49E | 464 ± 44 | 51.0 ± 1.41 |
| RadD R145E | 698 ± 75 | 28.4 ± 1.02 |

We next examined whether inclusion of SSB altered ATPase activity for WT RadD and the SSB site variants. As observed previously (17), titration of SSB from 0 to 40 μM increased the ATPase rate of WT RadD ∼3-fold (Figure 3C). In contrast, SSB failed to stimulate ATPase activity in the RadD variants even at the highest concentration tested. Thus, direct binding of RadD to SSB appears essential for SSB stimulation of RadD ATPase activity.

### SSB-binding pocket Arg mutants stimulate RecA-mediated strand exchange

RadD stimulates RecA-mediated strand exchange and function as a RecA accessory protein during recombinational repair (9). To test the importance of the RadD-SSB interaction in RecA mediated strand exchange stimulation, the activity of the SSB-binding RadD variants was tested using *in vitro* strand exchange reactions. Strand exchange reactions (Figure 4A) using a linear double-stranded (ds) DNA and complementary circular ssDNA were monitored for RadD stimulation. Control RecA-mediated strand exchange reactions with no RadD produced joint molecules (JM) and nicked circular (nc) DNA within five minutes of the start of the reactions (Figure 4B to D). As previously reported (9), the addition of WT RadD stimulated the RecA-mediated strand exchange determined by the presence of JM and nc DNA within two minutes of the start of the reactions (Figure 4B to D). Next, each charge-reversal RadD variant was tested for stimulation of RecA-mediated strand exchange. Each of the variants stimulated RecA-dependent strand exchange activity determined by the presence of JM and nc DNA within two minutes of the reaction start time (Figure 4b to D). Thus, alteration of the SSB-interacting pocket does not significantly alter the RecA stimulation functions of RadD.

**Figure 4:**
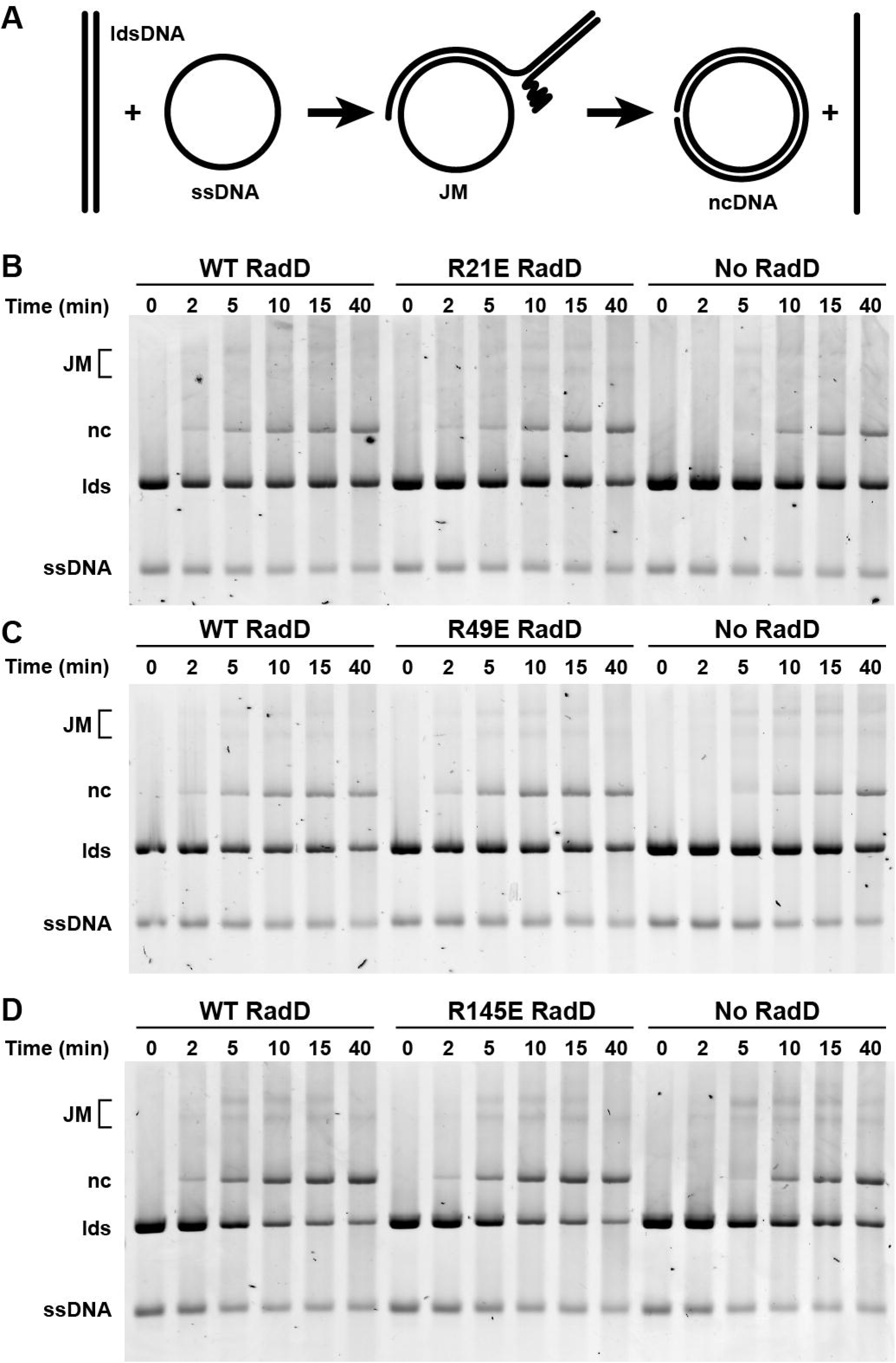
RecA-mediated strand exchange stimulation of RadD SSB-binding mutants. A) Schematic of *in vitro* strand exchange using linear dsDNA and a complimentary circular ssDNA. RecA-mediated strand exchange DNA progresses to joint molecules (JM) and eventually result in nicked circular (nc) and linear single stranded DNA products. B-D) Strand exchange mediated by RecA with either WT RadD, RadD SSB-binding variants, or no RadD. Reactions were conducted with RecA at a concentration of 6.7 μM and 6.7 nM wild type or variant RadD. Circular ssDNA, linear dsDNA (lds), nicked strand exchange product (nc), and joint molecules (JM) labeled. Experiment was performed 3 times, with representative data shown.

### SSB-binding radD mutants affect cell response to DNA damage

To examine the cellular effects of a loss of SSB binding by RadD, mutant *E. coli* strains were constructed in which the *radD* gene was mutated to encode the Arg21Glu, Arg49Glu or Arg145Glu RadD variants. Each of the *radD* mutant strains grew at wild-type rates and showed little to no increased sensitivity to DNA damaging agents relative to the control wild-type strain (Figure 5A, 6A, 6C, and S1A). SOS induction was measured using GFP expression under an SOS inducible promoter to gauge the effect of losing RadD-SSB interactions on stress signaling. GFP fluorescence indicates that, like *ΔradD* cells (8), SSB-binding *radD* mutants exhibited little change in SOS induction on their own (Figure S1B).

**Figure 5.**
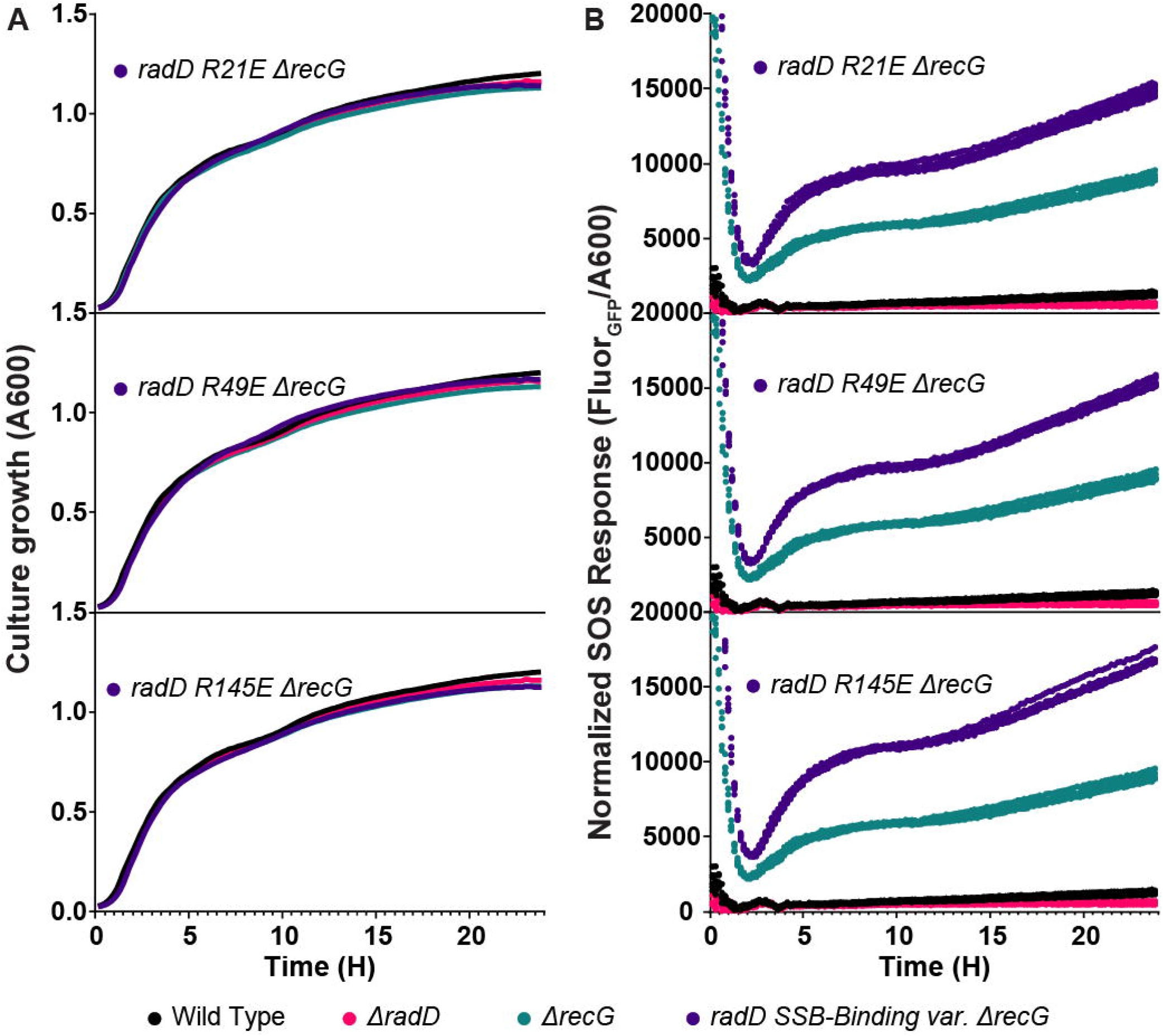
Effect of *radD* SSB-binding mutants on SOS response A) Liquid culture growth of *ΔrecG radD* SSB-binding mutant strains in comparison to wild type, *ΔrecG*, and *ΔradD*. Data points are the mean of 6 independent measurements. B) SOS response of *ΔradD, ΔrecG*, and *ΔrecG ΔradD* SSB-binding mutants measured using GFP fluorescence under *recN* promoter, normalized to A_600 nm_. Experiment was performed 6 times, with all data points plotted.

**Figure 6.**
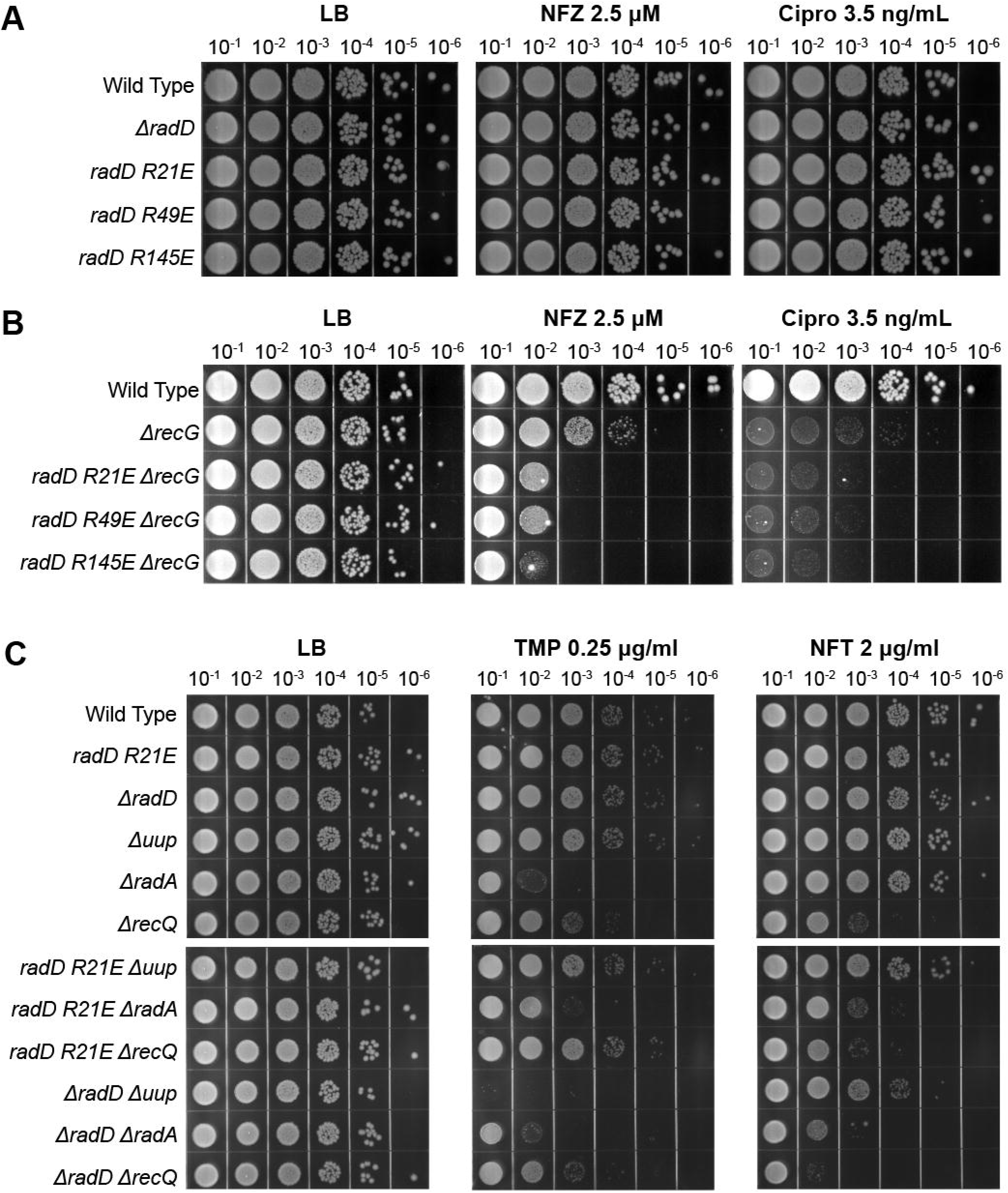
Drug induced DNA-damage stress sensitivity assays of chromosomal SSB-binding *radD* mutants in combination with DNA remodeling and repair enzymes. Each plate set is a record of drug treatment, dose, and serial dilution factor. A) Nitrofurazone and ciprofloxacin spot plates of SSB-binding mutants and *ΔradD* B) Nitrofurazone and ciprofloxacin spot plates of SSB-binding mutants in a *ΔrecG* background compared to single mutant *ΔrecG* strain. C) Sensitivity assays combining *radD* Arg21Glu chromosomal mutant with *uup, radD* and *recQ* deletions separately, compared to wild type *radD* or *ΔradD* conditions. Experiment was performed 3 times, with representative data shown.

Past studies have shown that *ΔradD ΔrecG* cells have severe growth defects and extreme sensitivity to DNA damaging agents (8). To enhance the possible effects of the RadD variants, *recG* deletions were introduced into each SSB-binding *radD* mutant strain. Unlike strains carrying deletions of both *radD* and *recG*, the resulting double mutants had no measurable growth defect in liquid media (Figure 5A). However, highly synergistic SOS induction was detected for all three *radD* SSB-binding mutants in the absence of *recG* (Figure 5B), indicative of increased DNA damage stress. The increase in SOS induction was detected in a low to no stress environment produced in normal nutrient-rich Luria Broth growth conditions. Thus, loss of interaction with SSB in the RadD variants does impact cellular function in cells lacking the RecG helicase but not to the extent of a full *radD* deletion.

Drug induced DNA-damage stress sensitivity assays of SSB-interaction *radD* mutants were carried out in the presence of gene deletions known to sensitize *ΔradD* cells to DNA damaging agents or previously linked to RadD function to better understand the function of the RadD-SSB interaction in a cellular context (8, 13–15). The compounds utilized in Figure 6 induce DNA damage through different modes of action. Nitrofurazone (NFZ) induces base lesions leading to ssDNA gaps through the formation of bulky deoxyguanosine adducts requiring nucleotide excision repair (38, 39). Ciprofloxacin (cipro) induces topological stress and dsDNA breaks through inhibition of DNA gyrase (40). The inhibition of DNA gyrase leads to a covalent replication block which devolves into ds breaks through replication fork collapse and protein degradation (41, 42). Trimethoprim (TMP) inhibits thymine synthesis which cascades to nucleotide depletion, replisome stalling and eventual DNA damage (43). Nitrofurantoin (NFT) functions by activation through bacterial nitroreductase causing metabolic stress through activated nitrofurantoin byproducts (44). These byproducts cause a broad range of damage to protein enzymes, RNA and DNA. The resulting situations and structures challenge DNA repair and cause total protein synthesis inhibition at high nitrofurantoin concentrations (45–47).

Each *radD* mutant was first tested for sensitivity to NFZ and cipro in a *ΔrecG* background. Notably, the *ΔradD ΔrecG* double mutant (including a suppressor mutation in *priA* that permitted growth) had previously been shown to be sensitive to both DNA damaging agents (8). Single point *radD* SSB-binding mutants and *ΔradD* strains were not sensitive to NFZ and cipro at the assayed drug concentrations (Figure 6A). However, all three SSB-interaction *radD ΔrecG* mutants displayed increased sensitivity to low concentrations of NFZ and cipro (Figure 6B). At the assayed concentration of NFZ (2.5 μM), *ΔrecG* cells show ∼2-log sensitivity compared to wildtype *E. coli* strain. The three SSB-interaction *radD ΔrecG* mutants had an additional increase in nitrofurazone sensitivity of about 2-3 logs when compared to *ΔrecG* cells. Cells bearing a *recG* deletion have a lower plating efficiency and impaired growth with cipro, leading to faint patterns in sensitivity assays. Even so, all three *radD* SSB-binding *ΔrecG* mutants have a lower plating efficiency compared to *ΔrecG* cells (Figure 6B). The increase in basal SOS induction in high nutrient liquid culture and the sensitivity of *radD* SSB-binding and *ΔrecG* double mutant cells to DNA damaging agents indicates that RadD interaction with SSB is important for RadD function during DNA repair *in vivo*. However, deficiencies in RadD SSB-binding are insufficient to fully abolish RadD function since *radD* SSB-binding mutants do not exhibit a full *radD* deletion phenotype. This suggests that the RadD-SSB interaction is important in pathways that complement the function of RecG in post-replication gap repair and dsDNA break repair.

Lastly, Arg21Glu *radD* strains were tested for sensitivity in the absence of other DNA repair genes previously reported to sensitize cells in a *ΔradD* context (8, 13–15). Specifically, deletions of *uup, radA*, and *recQ* were tested in combination of Arg21Glu *radD* and *ΔradD*. TMP and NFT were utilized as DNA damaging agents as *ΔradD Δuup* and *ΔradD ΔrecQ* cells are known, respectively, to be sensitive to these damaging agents (8, 13). Of the three SSB binding mutants of *radD*, only the *radD* Arg21Glu mutation was used moving forward as no difference in phenotypes was observed in earlier experiments between *radD* Arg21Glu, *radD* Arg49Glu, *and radD* Arg145Glu.

The *ΔradA* mutant exhibited little to no sensitivity to NFT, while the *ΔradD ΔradA* strain exhibited a severe sensitivity phenotype of -3 log difference. The *radD* Arg21Glu *ΔradA* strains exhibited an intermediate phenotype between wild-type and a full deletion of *radD* (Figure 6C). Unexpectedly, the TMP sensitivity of *ΔradA* cells is slightly alleviated by the presence of the Arg21Glu SSB-binding mutant of *radD*. The marked effect of decreased sensitivity to TMP is not observed in the double deletion *ΔradD ΔradA* strain. TMP causes stress via a decline in nucleotide concentration, which may increase formation of ssDNA gaps as polymerases stall and restart in other locations. The effects seen with TMP indicate that RadD is probably an important part of at least one pathway directed at repairing these gaps, but the RadD-SSB interaction is not important and may perhaps be a bit deleterious in this role.

Similar patterns are observed in *radD* SSB-binding mutants treated with TMP in a *ΔrecQ* background. However, the presence of an SSB-interaction mutation does not affect the sensitivity of *ΔrecQ* cells to NFT. Both *ΔrecQ* deletion and *radD* Arg21Glu *ΔrecQ* mutants have about a -3 log difference in sensitivity to NFT compared to wild-type cells, while *ΔradD ΔrecQ* cells have a -4 log difference in viability when exposed to NFT. The added sensitivity of *ΔradD ΔrecQ* cells to NFT is a new finding, which further cements the function of RadD in the remodeling of DNA intermediates as deletion of *radD* affects *recQ* deficient cells. However, this newly detected genetic interaction is not affected by the absence of RadD-SSB interactions.

Out of all known *radD* genetic interaction mutants, *Δuup* cells were the least affected by the *radD* SSB-interaction mutation, leading to no discernable difference in sensitivity of *radD* Arg21Glu *Δuup* compared to *Δuup*. While this was the case, the double mutant *ΔradD Δuup* strain was still sensitive to NFT and extremely sensitive to TMP (Figure 6C).

Overall, charge reversal of residues important for SSB-interaction in *radD* lead to increased sensitivity to certain DNA damaging agents in strains deficient for additional DNA metabolism genes. This demonstates the importance of the SSB-RadD interaction *in vivo*, while also highlighting a separation of function in the *radD* SSB-interaction mutants leading to possible split pathways to the known function of RadD in DNA repair.

## Discussion

The work in this study leads to four conclusions. First, we have identified the SSB binding pocket on the RadD protein. The pocket is defined by a hydrophobic pocket framed by Arg residues 21, 49, and 145. Second, charge-reversal mutations of Arg residues surrounding the binding pocked eliminate RadD-SSB interaction *in vitro* without significantly altering the RadD ATPase activity in the absence of SSB. However, SSB fails to stimulate ATPase activity in the variants. Third, the elimination of the RadD-SSB interaction has measurable effects *in vivo. radD* mutants encoding variants with impaired SSB-binding properties grow as well as wild-type *E. coli* under normal conditions. However, the mutants exhibit a synergistic increase in DNA damage sensitivity when paired with deletions of *radA* and *recG*. Fourth, the phenotypes of the SSB-binding mutants of *radD* are significant but not as severe as those exhibited by *radD* deletion strains. Thus, there appear to be some RadD functions that rely on interaction with SSB and others that do not. RadD-SSB appears to be important for recombinational processes that are part of post-replication gap repair and dsDNA break repair. The previously identified activity of RadD in stimulating RecA-mediated DNA strand exchange does not depend on an SSB interaction, at least *in vitro*. In repairing gaps created by TMP, the RadD-SSB interaction appears less important and perhaps even slightly deleterious.

The identified SSB-binding pocket of RadD was predicted to bind the C-terminal tail of SSB through interactions with three key Arg residues (21, 49, and 145). Typical SSB C-terminus interactions are coordinated by either Arg or Lys side chains interacting with the terminal α-carboxyl group of SSB and with negatively-charged Asp side chains within the SSB-Ct (18–26). Charge reversal mutagenesis of Arg21, Arg49, or Arg145 results in a severe reduction in binding to SSB, which is typical of the results seen for other SSB-interacting proteins and their SSB interaction pockets (19–28). Moreover, ATPase activity in the RadD variants was no longer stimulated by SSB, indicating that binding at the SSB-binding stie was critical for stimulation. How might SSB binding induce ATPase activity of RadD? To answer this question the structure of RadD bound to ADP (16) was compared to the top RadD/SSB-Ct model, aligning the two models using their RD1 domains. The domains align with an RMSD of 1.45 Å for all α-carbons with most catalytic motifs in the aligned structure overlaying one another. However, a helix linking the SSB binding site and the P-loop (motif I) in RadD was shifted, adjusting the position of an essential catalytic Lys by ∼2 Å within the ATPase active site (Figure 7). As noted in the prior RadD/ADP crystal structure (16), motif I is not properly aligned in the structure to support ATPase function. It may be that SSB binding to RadD at one end of the linker helix alters the position of the motif I at the other end, making the active site more optimal for ATPase activity. This is similar to an allosteric stimulation mechanism that has been proposed for the RecQ DNA helicase in which DNA binding at one site alters the position of a helix connected with motif II to improve the position of the motif in the ATPase active site (34).

**Figure 7.**
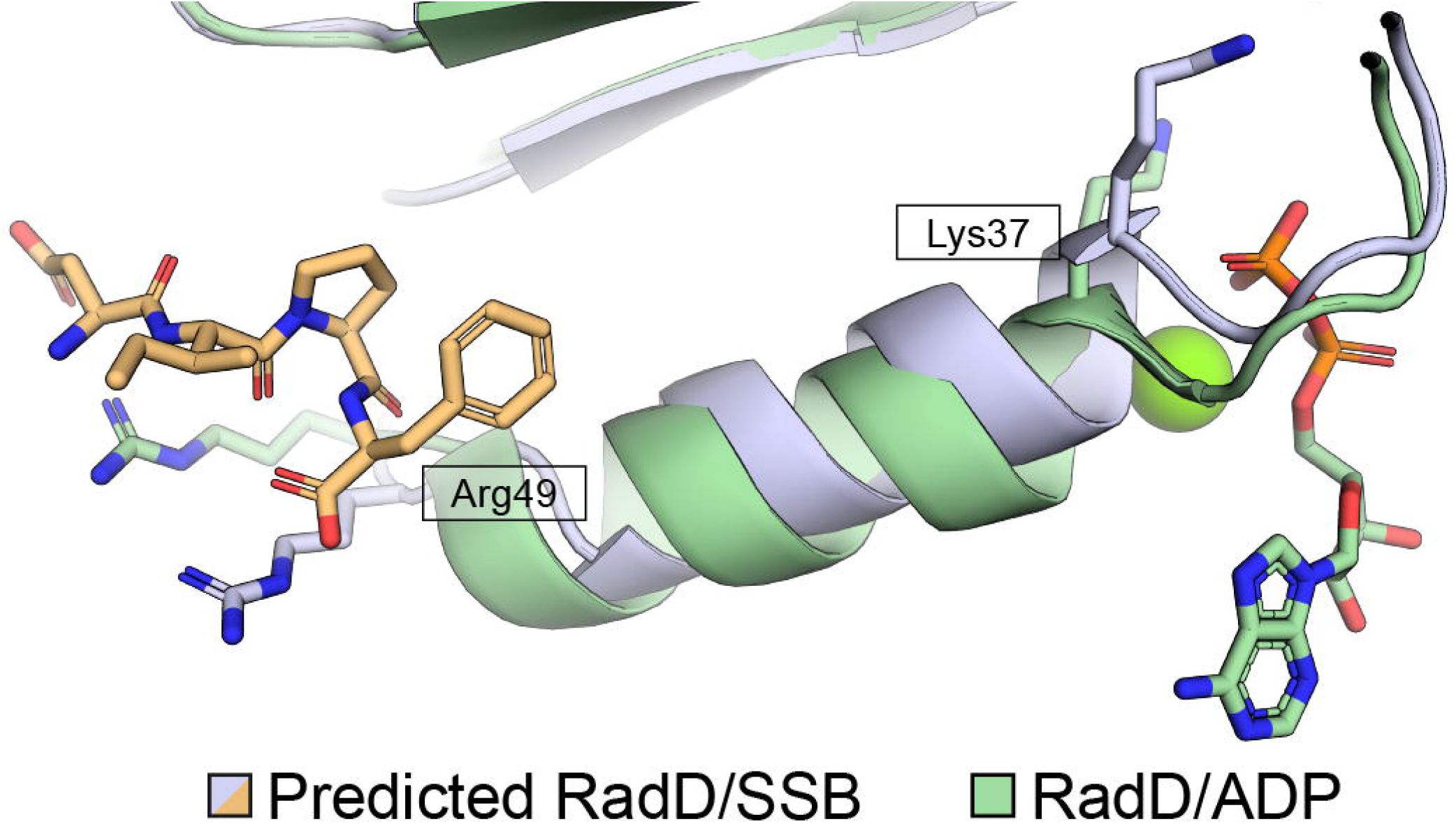
Predicted SSB-mediated RadD ATPase induction. RadD <u>R</u>ecA-Like <u>d</u>omain <u>1</u> (RD1) alpha carbon alignment of predicted RadD-SSB interaction (grey) to experimentally derived RadD-ADP complex (green) (PDB 7R7J (16)) with an RMSD of 1.4 Å. Positional differences for Arg49 and Lys37 are notable in comparing SSB-bound and unbound state using the AlphaFold predicted RadD/SSB C-terminus structure and published RadD/ADP structure.

Mutations of arginines important for SSB-interactions in *radD* lead to increased SOS signaling and sensitivity to DNA damaging agents in the absence of repair genes known to function along with *radD* (9, 13–15). The effect of the SSB-binding mutations varied depending on the type of DNA damaging drug used as well as the deletion background. This differentiation allows for a direct evaluation of the importance of the SSB interaction for *in vivo* RadD function in different pathways.

The most severe effect was observed with cells with *radD* mutations and a *recG* deletion (8). This finding, along with the fact that both RadD and RecG function as repair intermediate processors posits the hypothesis that RadD and RecG are vital to the cell through DNA repair intermediate resolution (8, 9, 11, 48, 49). Repair intermediates are a constant in dividing cells due to the ever-present need for gap repair and require resolution, as prolonged presence of branched intermediates is toxic (8). The extreme sensitivity of *ΔradD ΔrecG* mutants suggest both genes function in overlapping but distinct manners vital to cell survival. Abrogation of the SSB-interaction leads to an intermediate phenotype in *ΔrecG* cells during induced dsDNA breaks or DNA gap formation stress, indicating that the SSB interaction is important for the overall activity of RadD. However, the *radD* SSB-interaction mutations led to no noticeable change in sensitivity to cells bearing a *uup* deletion under DNA replicative stress. Uup is predicted to function upstream in the RecG pathway, binding and remodeling Holliday Junction structures (8). This indicates that RadD SSB-binding activity does not play a role in the interplay of *radD* and *uup* in suppressing DNA crossover events. The current results indicate a separation of SSB-dependent and SSB-independent functions of RadD in DNA repair.

RadD was also recently shown to stimulate RecA dependent DNA strand exchange *in vitro* (9). However, abrogation of RadD-SSB interactions did not affect the function of RadD as a RecA-dependent strand exchange accessory protein (Figure 4). These findings leave a discrepancy between the *in vivo* importance of RadD-SSB interactions and missing *in vitro* effect on the only known molecular activity of RadD. One explanation for these findings is that the importance of SSB-RadD interactions may differ in the full context of *in vivo* strand exchange not captured by *in vitro* assays. Furthermore, RadD is important for both RecA-dependent (9) and RecA-independent (13) branch intermediate processing, which provides a second explanation for the differing requirements for RadD-SSB interactions. There is a possibility of the RadD-SSB interaction being exclusively important to RecA-independent pathways.

Questions remain as to the function of the RadD-SSB interaction in RadD repair pathways, particularly about the role that SSB stimulation plays in branch intermediate processing and the possibility of separate activity of RadD in RecA-dependent and independent DNA repair pathways. Further experiments are required to build a model of RadD dependence on SSB interaction on differing DNA repair pathways. This work has made such future research possible by characterizing the SSB-RadD interaction through a pocket in RadD.

## Experimental procedures

### Alphafold2 model of the RadD-SSB C-term interaction

A model of the RadD-SSB-Ct interaction was created using ColabFold software (31), which incorporates MMseqs2 (50) sequence alignment and the AlphaFold2 program (30) to predict the structure of multimer complexes. RadD and SSB C-terminal peptide (Asp-Ile-Pro-Phe) sequences were input to ColabFold and run using default settings (multiple sequence alignment = MMSeqs2 (UniRef+Environmental), pair_mode = unpaired+paired, model type= auto, number of cycles = 3). The top scoring model was taken to build the RadD-SSB model.

### Evolutionary conservation of RadD pocket

ConSurf software (51, 52) was used to align 200 RadD sequences across bacterial species with a maximal percent identity of 95% and minimal percent identity of 50% to the ADP-bound RadD crystal structure (16). Homologs were collected from UNIREF90 and Multiple Sequence Alignment was built using MAFFT. A HMMER homolog search algorithm was used with an E-value of 0.0001. Resulting conservation scores were visualized using PyMOL (53).

### Purification of RadD variants

Individual *E. coli radD* mutants (Arg21Glu, Arg49Glu and Arg145Glu) were created for overexpression by *in vivo* assembly cloning (54) using WT *radD* overexpression plasmid pEAW724 (17) and resulting constructs were verified by sequencing. *E. coli* STL2669/pT7pol26 cells were transformed with pEAW724 or plasmids encoding RadD variants and grown at 37°C in Luria Broth supplemented with 100 μg/ml ampicillin and 40 μg/ml kanamycin. Cells were induced with 1 mM isopropyl β-D-thiogalactopyranoside at mid log phase (A_600_ ∼0.6 A) and grown for an additional 3 hours. Pelleted cells were resuspended in lysis buffer (25% (w/v) sucrose, 250 mM Tris chloride (pH 7.7), 7 mM EDTA, 1 μM pepstatin, 1 μM leupeptin and 0.1 mM phenylmethylsulfonyl fluoride) and lysed by the addition of 5mg/ml lysozyme and sonication on ice. Lysate was clarified by centrifugation and RadD (or variants) was precipitated from the soluble fraction by slowly adding solid (NH_4_)_2_SO_4_. Protein pellets were then resuspended in R-buffer (20 mM Tris chloride (pH 7.7), 1 mM EDTA, 10% glycerol) + 1 M (NH_4_)_2_SO_4_ and loaded on to a butyl-Sepharose column. RadD was eluted in R buffer with a gradient of one to zero molar (NH_4_)_2_SO_4_ over five column volumes. Fractions containing RadD were pooled and dialyzed against a buffer containing 20 mM phosphate (pH 7.0), 200 mM KCl, 1 mM EDTA, and 10% glycerol, loaded on a ceramic hydroxyapatite column, and collected in the wash. RadD was then dialyzed into R-buffer + 200 mM KCl and subsequently run over both Source-15S and Source-15Q columns. RadD and variants flowed through while the remaining contaminants bound to either of the ion exchange columns. Purified RadD was then concentrated, flash frozen and stored at -80°C. The purified protein was >95% pure by gel and free of any detectable nuclease activity. Concentration was determined using an ε _280_ of 5.59 × 10^4^ M^−1^cm^−1^ (17) for the wildtype and RadD variants.

### RecA purification

RecA was purified as previously described (55) and the concentration determined utilizing an ε_280_ of 2.23 × 10^4^ M^-1^ cm^-1^ (56).

### SSB purification

Full length SSB was purified as previously described (57) and the concentration determined utilizing an ε_280_ of 2.38 × 10^4^ M^−1^cm^−1^. Fluorescein amide-labeled SSB C-terminal peptide (5-FAM WMDPDDDIPF) was synthesized and purified commercially (GenScript).

### Fluorescence anisotropy

Increasing concentrations of Rad or variant were incubated with 10 nM 5-FAM SSB C-terminal peptide in a reaction buffer composed of 25 mM Tris-acetate (pH 7.5), 200 mM potassium glutamate, 10 mM magnesium acetate, 1 mM DTT, and 0.1 μg/ml bovine serum albumin for 15 minutes at room temperature. Fluorescence anisotropy of triplicate samples were immediately measured using a Beacon 2000 fluorescence polarization system with an excitation and emission wavelengths of 490 nm and 535 nm, respectively. Data were plotted in GraphPad prism 9.4.1 and fit curves were generated using a one-site nonlinear regression model.

### Ammonium sulfate co-precipitation

Co-precipitation experiments were performed as described previously (32). Pellet fractions were resuspended in 30 µL of loading buffer and run on a 12% SDS-PAGE gel.

### ATPase reactions

ATP hydrolysis was monitored using a coupled spectrophotometric enzyme assay (58) using increasing concentrations of SSB with a constant concentration of RadD or RadD variant. A Varian Cary 300 UV-Vis Bio Spectrophotometer equipped with a temperature controller was used to measure NADH oxidation at 380 nm coupled to an ATP regeneration system in triplicate. The reactions were carried out at 37 °C in 25 mM Tris-acetate (80% cation, pH 7.5), 200 mM potassium glutamate, 10 mM magnesium acetate, 1 mM DTT, 5 % (w/v) glycerol, an ATP regeneration system (10 units/ml pyruvate kinase, 2.2 mM phosphoenolpyruvate), a coupling system (2 mM NADH and 10 units/ml lactate dehydrogenase), and 200 nM purified WT RadD or variants. ATP concentration was 3 mM unless otherwise stated.

### Strand exchange reactions

All strand exchange reactions were caried out at 37°C. 20 μM (nt) ΦX174 Virion DNA (New England Biolabs) was incubated in 1X RecA buffer (25 mM Tris-acetate (80% cation, pH 7.5), 5% (w/v) glycerol, 3 mM potassium glutamate, and 10 mM magnesium acetate), 1 mM DTT, 2.5 mM phospho(enol)pyruvate (PEP), 10 U/ml pyruvate kinase, and 6.7 μM RecA for 10 minutes. The reactions were incubated for another 10 minutes after the addition of 2.1 μM SSB and 3 mM ATP. Each reaction was initiated by the addition of 20 μM nt ΦX174 RF1 DNA (New England Biolabs) previously digested by PstI, and 6.7 nM RadD or variant. Aliquots (10 μL) were taken at each time point and quenched for 10 minutes at 37°C with 5 μL of 3:2 6× Ficoll:10% SDS. Samples were then run on an 0.8% TBE agarose gel at 25 V for 16 hrs.

### Chromosomal mutant strain construction

Strains with gene deletions or mutations in their native loci were created by a modified version of the Datsenko and Wanner (59) method. pEAW507, which includes a mutant FRT-Kan^R^-wt FRT cassette, was PCR amplified with primers containing of 21 nucleotides of homology with the ends of the cassette and 50 nucleotide sequences flanking the loci of interest. The resulting PCR product was gel purified and electroporated into cells containing pKD46. The pKD46 plasmid is an expression vector for λ Red recombinase containing ampicillin resistance. Recombinase expression and subsequent reaction was induced using L-arabinose. Plasmid cured colonies with kanamycin resistance and ampicillin sensitivity were screened for gene deletions through PCR confirmation and sequencing.

A similar approach was used to create chromosomal mutants. The mutant gene was cloned into pET21 followed by the kanamycin FRT cassette resulting in a plasmid containing the mutant gene-mutant FRT-Kan^R^-wt FRT cassette. This was then used as template to PCR amplify and recombine into MG1655 as previously described.

### SOS response

SOS induction was monitored using a plasmid-based GFP reporter assay. A plasmid (pEAW903) expressing SuperGlo GFP under the control of the SOS inducible *recN* promoter was transformed into target strains. Overnight transformant cultures were diluted 1:100 in fresh Luria Broth and transferred to a black-walled, clear-bottomed 96 well plate in sextuplicate. Culture growth and GFP fluorescence were monitored at 600 nm and 488/515 nm every 10 minutes while orbital shaking at 37°C for 24 hours using an H1 Synergy Biotek plate reader. GFP fluorescence was normalized to culture growth, resulting in the SOS induction curves. SOS induction curve points prior to ∼2 hours are exaggerated due to background from overnight growth fluorescence and low OD readings leading to large uncertainty and should be ignored.

### DNA damage sensitivity plating

Overnight cultures were used to inculcate 5 ml of fresh Luria Broth in a 1:100 ratio and grown to an OD_600_ of 0.2 at 37°C. Cultures were serially diluted (10^−1^ to 10^−6^) in 1xPBS buffer on a 96 well plate. Culture dilutions were then spot plated on Luria Broth agar plates made with the indicated DNA damaging agent concentration. Plates were grown at 37°C overnight and imaged the next day.

## Supporting information

This article contains supporting information.

## Funding and additional information

This work was supported by grant RM1 GM130450 from the National Institute of General Medical Sciences. The content is solely the responsibility of the authors and does not necessarily represent the official views of the National Institutes of Health.

## Abbreviations

Cipro: The abbreviations used are
dsDNA: ciprofloxacin
DNA: double stranded
FAM: fluorescein amidite
JM: joint molecules
ncDNA: nicked circular DNA
NFT: nitrofurantoin
NFZ: nitrofurazone
RD1: RecA-like domain 1
RD2: RecA-like domain
SF2: helicase, superfamily 2 helicase
SSB: single-stranded DNA-binding protein
SSB-Ct: SSB C-terminus
ssDNA: single stranded DNA
TMP: trimethoprim
WT: wildtype.

