## Supplemental Figure 1 for "Interaction with single-stranded DNA-binding protein modulates *Escherichia coli* RadD DNA repair activities"

**Supporting Information**

**Figure S1**

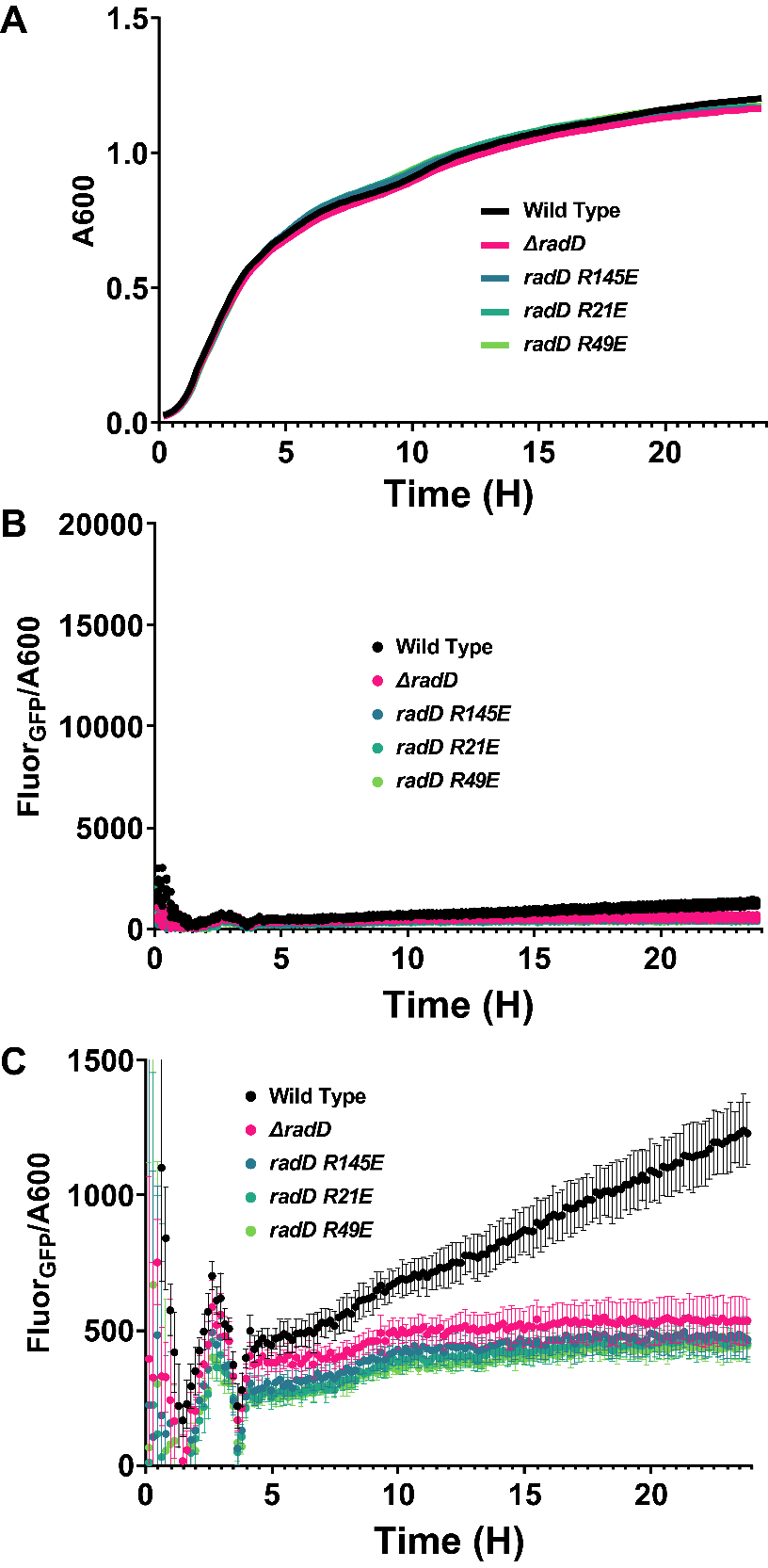

**Figure S1** *SOS induction of radD SSB-binding mutants* A) Growth curves of *ΔradD* and SSB-binding *radD* mutant strains compared to wildtype *E. coli*, all transformed with SOS inducible GFP plasmid. Data points are the mean of 6 independent measurements. B) Normalized SOS induction measured by GFP fluorescence induced by P_recN_ of wildtype, *ΔradD* and radD SSB-binding mutant strains. Using the same Y-axis scale as Figure 4. Data points are the mean of 6 independent measurements. C) Same data as B with a zoomed in Y-axis scale to highlight differences in the SOS curves. Data points are the mean of 6 independent measurements with error bars representing the standard error.
